# Lipid Transfer Proteins and PI4KIIα Generate a Phosphoinositide-Linked Proteome

**DOI:** 10.64898/2025.12.17.694959

**Authors:** Noah D. Carrillo, Mo Chen, Poorwa Awasthi, Tianmu Wen, Xiangqin Chen, Giang Vu, Dhruv Brahmbhatt, Michael Herlihy, Benjamin B. Minkoff, Michael Sussman, Vincent L. Cryns, Richard A. Anderson

## Abstract

Phosphoinositide (PIP_n_) lipid second messengers in membranes regulate a myriad of cellular processes. In the cytosol, the phosphatidylinositol (PI) 3-kinase (PI3K)/Akt pathway is scaffolded on IQGAP1 to facilitate the activation of Akt by the synthesis of PI3,4,5P_3_. In the nucleus, PIP_n_ signaling occurs in regions devoid of membranes via their stable association with proteins. While several of these proteins have been identified, understanding the extent and impact of protein-linked PIP_n_ signaling warrants further investigation. The tumor suppressor p53, was shown in the companion paper to be regulated by PI transfer proteins (PITPs) and a PI 4-kinase (PI4KIIα), which are required to form p53-PIP_n_ complexes that assemble a nuclear PI3K/Akt pathway. Here we report that class I PITPs (PITPα/β) and PI 4-kinase initiate PIP_n_ linkages to many different proteins. PITPα/β and PI4KIIα accumulate in the nucleoplasm in response to stress and are necessary to synthesize nuclear PIP_n_s linked to proteins. These PITPα/β-dependent protein-PIP_n_ complexes are detected by metabolically labeling cells with the PIP_n_ precursor [H^3^]-*myo*-inositol and resist denaturation and SDS-PAGE, indicating that these protein-PIP_n_ complexes represent a putative posttranslational modification. Proteomic analyses of proteins that are regulated by PITPα/β and/or are linked to PI4,5P_2_ have identified an emerging PIPylome that is enriched in metabolic, signaling, cytoskeletal and DNA repair pathway components. Taken together, these data provide evidence for an emerging proteome with linked PIP_n_s that represent a PIP_n_ signaling paradigm that is distinct from the membrane-localized pathway but utilizes many of the same PIP kinases and phosphatases.

**In brief:** Phosphatidylinositol transfer proteins and PI 4-kinase initiate a PIP_n_-linked protein network in membrane-free regions.

## Introduction

Phosphoinositide (PIP_n_) lipid second messengers are synthesized in membranes from phosphatidylinositol (PI) by PIP kinases and phosphatases to regulate metabolism, membrane trafficking, motility, cell growth and death^1–5^. Strikingly, the PI 3-kinase (PI3K)/Akt pathway is scaffolded on IQGAP1, which uses PI to synthesize PI3,4,5P_3_ by an ordered assembly of PIP kinases ending with the PI3K generation of PI3,4,5P_3_ that recruits and activates the serine-threonine kinase Akt pathway on membranes^6, 7^. PIP_n_ signaling also occurs in non-membranous regions of the nucleus and these signaling competent PIP_n_s are resistant to detergent extraction, indicating a stable interaction^8–12^, that appears distinct from conventional lipid compartments^4, 13–17^. Consistent with this non-canonical PIP_n_ signaling, six of the seven PIP_n_ isomers (all but PI3,5P_2_), as well as PIP kinases and phosphatases, have been reported in non-membranous cellular compartments in the nucleus^4, 9, 18^. Several noteworthy PIP_n_-linked proteins have recently been identified including murine double minute 2 (MDM2)^19^, nuclear speckle targeted PIPKIα regulated-poly(A) polymerase (Star-PAP)^20^, Yes-associated protein (YAP) and transcriptional coactivator with PDZ-binding motif (TAZ)^16^, nuclear factor erythroid 2-related factor 2 (NRF2)^21^, and the tumor suppressor p53^14^. In each case, PIP_n_-linkage is resistant to denaturation/SDS-PAGE and is metabolically labelled with the PI precursor [^3^H]-*myo*-inositol. Furthermore, small heat shock proteins such as αB-crystallin and Hsp27 are recruited to PIP_n_-linked proteins and regulate protein stability and likely other functions. For p53, similar to of IQGAP1, the regulated assembly/disassembly of p53-PI3,4,5P_3_ by PIP kinases and PTEN, respectively, controls Akt activation in the nucleus and may be a mechanism for mutant p53’s oncogenic activity^13^.

In the companion paper to this manuscript^22^, we discovered novel regulators of the p53-PI3K/Akt pathway, namely, the class I (α/β) PI transfer proteins (PITPs)^23, 24^ and the PI 4-kinase PI4KIIα^25^, which were previously thought to function only in the cytosol. Specifically, we demonstrated that PITPα/β and PI4KIIα interact with p53 in the nucleus and initiate the synthesis of p53-PIP_n_ complexes^22^. Given the existence of other PIP_n_-linked proteins, we postulated that PITPα/β and PI4KIIα might be key components of a broader program to link PIP_n_s to other proteins. Here we show that this is indeed the case. PITPα/β and PI4KIIα are required to initiate stable PIP_n_ linkages to a collection of cellular protein (coined the PIPylome) detected by immunoblotting and metabolic labeling of cells with the PIP_n_ precursor [^3^H]-*myo*-inositol. PI4,5P_2_ linked proteins were affinity purified and proteomic analyses was used to identify a PIPylome, providing additional evidence for a novel protein-linked PIP_n_ signaling paradigm.

## Results

### The nuclear localization of PITPs is regulated by cellular stress

The discovery that PITPα/β initiate nuclear p53-PIP_n_ signaling in the companion paper suggests that PITPs may function beyond their established roles in the cytosol^22^. To investigate this possibility, we first examined the subcellular localization of the smallest and most homologous members within the PITP family, class I PITPα and PITPβ and class II PITPNC1^26^ by immunofluorescence (IF). Each of these PITPs and PI4,5P_2_ were expressed in the nucleus under basal conditions, and the nuclear levels of each isoform and PI4,5P_2_ were increased by the DNA damage (Fig. 1a,b). The nuclear translocation of PITPα/β in response to genotoxic stress was observed across all cell lines examined (Fig. 1c and Extended Data Fig. 1a). Moreover, the nuclear levels of PITPs correlated with nuclear levels of PI4,5P_2_ and over 60% of the IF signal for PITPα/β was in the nucleus after cisplatin treatment (Extended Data Fig. 1b,c). The basal and genotoxic stress-induced nuclear localization of PITPα/β was confirmed by co-IF with lamin A/C to define the nuclear envelope (Fig. 1d and Extended Data Fig. 2a-d). This was further validated using 3D sectioning (Extended Data Fig. 3a-d). Notably, multiple cellular stressors enhanced the nuclear localization of PITPα/β (Fig. 1e,f and Extended Data Fig. 3e). Taken together, these findings underscore that stress enhances the nuclear localization of PITPα/β.

**Figure 1.**
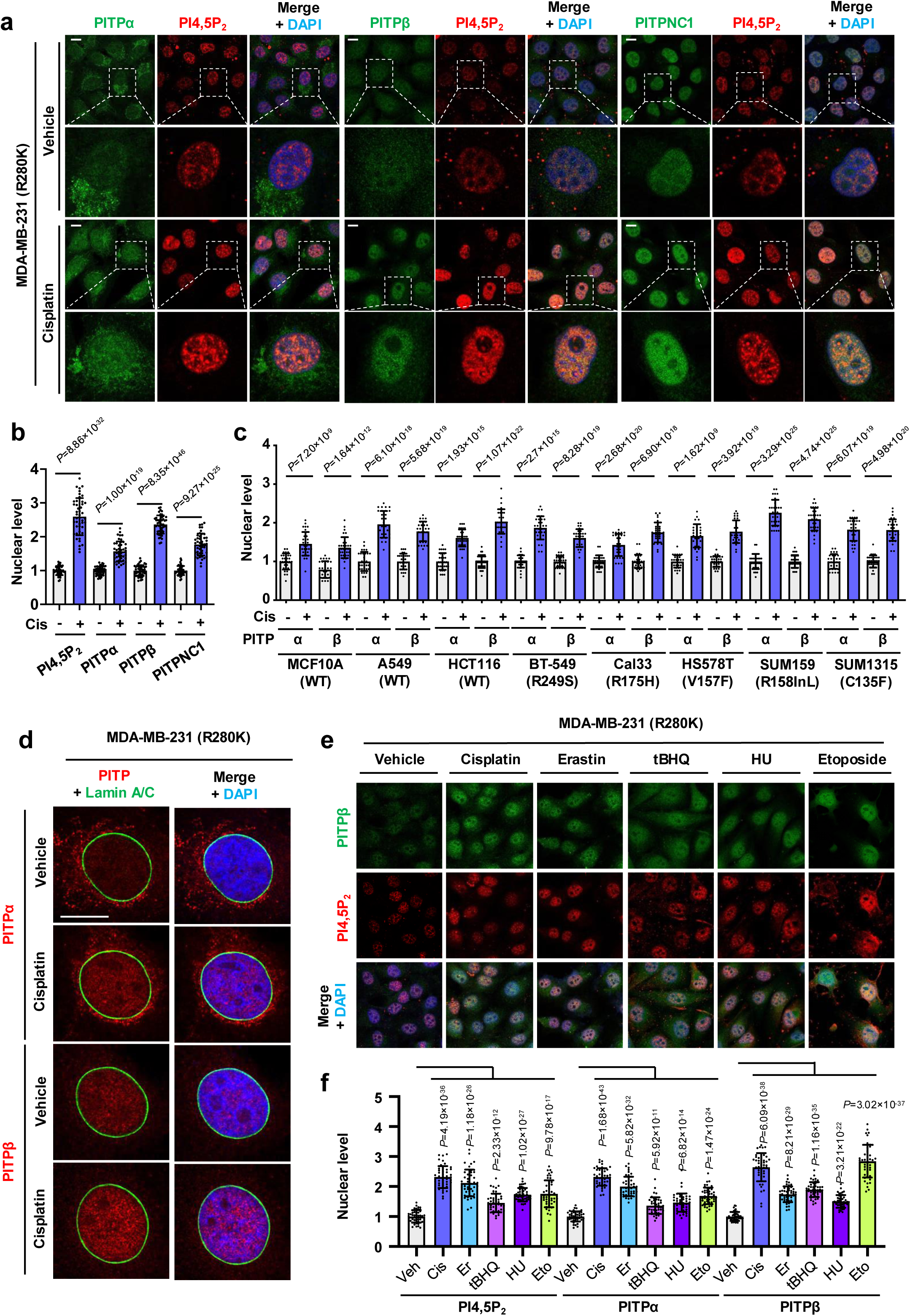
PITPα and PITPβ accumulate in the nucleus in response to stress. **a-b**, Confocal images of IF staining against PITPα/PITPβ/PITPNC1 and PI4,5P_2_ in MDA-MB-231 cells treated with vehicle or 30 µM cisplatin for 24 h. The nuclei were counterstained by DAPI. The nuclear levels of PITPs and PI4,5P_2_ were quantified by ImageJ (**b**). n=3, 15 cells from each independent experiment. *p* value denotes two-sided paired t-test. **c**, Quantification of IF staining against PITPα/β in wild-type and mutant p53 expressing cells, including MCF10A (WT), A549 (WT), HCT116 (WT), BT-549 (R249S), Cal33 (R175H), HS578T (V157F), SUM159 (R158InL), and SUM1315 (C135F). Nuclear PITPα and PITPβ levels were quantified by ImageJ. *p* value denotes two-sided paired t-test. See expanded images in Extended Data Fig. 1a. n=3 independent experiments. **d**, Confocal images of IF staining against PITPα or PITPβ overlaid with the nuclear envelope marker Lamin A/C in MDA-MB-231 cells treated with vehicle or 30 µM cisplatin for 24 h. The nuclei were counterstained by DAPI. n=3 independent experiments. **e-f,** MDA-MB-231 cells treated with 30 µM cisplatin, 20 µM erastin, 100 µM tBHQ, 100 µM etoposide, 100 µM hydroxyurea, or vehicle for 24 h. The cells were processed for IF staining against PITPα/β and PI4,5P2. The nuclei were counterstained by DAPI. The nuclear levels of PITPs and PI4,5P2 were quantified by ImageJ (**f**). See expanded images in Extended Data Fig. 3e. n=3, 15 cells from each independent experiment.

### PITPs regulate the nuclear PIP_n_ pool

A major enigma in the nuclear phosphoinositide field is the molecular and cellular characterization of PIP_n_ pools in non-membranous compartments^8–10^. PI4,5P_2_ detected by specific monoclonal antibodies form distinctive nuclear foci that colocalize with nuclear speckle markers in regions devoid of membranes and are increased in response to genotoxic stress^4, 8, 13, 14, 27^. These findings are reminiscent of the stress-enhanced nuclear localization of PITPα/β that occurs with increased nuclear PI4,5P_2_ (Fig. 1 and Extended Data Fig. 2a-d). Given that PI is the precursor of all PIP_n_s^3^, we investigated the role of PITPs in regulating nuclear PIP_n_ pools. Silencing PITPα or PITPβ reduced basal and stress-induced nuclear PI4,5P_2_ levels, while combined knockdown (KD) of both PITPα and PITPβ further decreased nuclear PI4,5P_2_ levels compared to the individual KD (Fig. 2a,b and Extended Data Fig. 4a). In contrast, PITPNC1 KD had little effect on nuclear PI4,5P_2_ levels (Fig. 2a,b). Consistent with these results, PITPβ knockout (KO) cell line was created from MDA-MB-231^Cas9^ cells. The combination of PITPβ KO and KD of PITPα abrogated PITPα/β IF in these cells validating antibody specificity (Extended Data Fig. 4b,c). Combined KD of PITPα and PITPβ also suppressed stress-induced nuclear levels of PI4P and PI3,4,5P_3_, although these PIP_n_s are largely cytoplasmic under basal nor stressed conditions (Fig. 2c,d). Combined KD of PITPα and PITPβ robustly inhibited stress-induced nuclear PI4,5P_2_ generation in a panel of human transformed and untransformed cells independent of p53 mutational status (Fig. 2e-h and Extended Data Fig. 4d,e). The KD of PITPα and PITPβ also reduced the levels for PI4P and PI3,4,5P3 detection, although these isomers where more cytosolic as previously reported ^28–32^. Moreover, the reduction in the stress-induced PI4,5P_2_ levels by combined PITPα and PITPβ KD using distinct siRNAs targeting their 3’-UTR was rescued by expression of PITPα^wt^, but not the PITPα^T59D^ mutant, which has diminished PI binding (Extended Data Fig. 5a-d). These results demonstrate that PITPα/β are necessary to generate nuclear PIP_n_ pools by a mechanism that requires PI binding.

**Figure 2.**
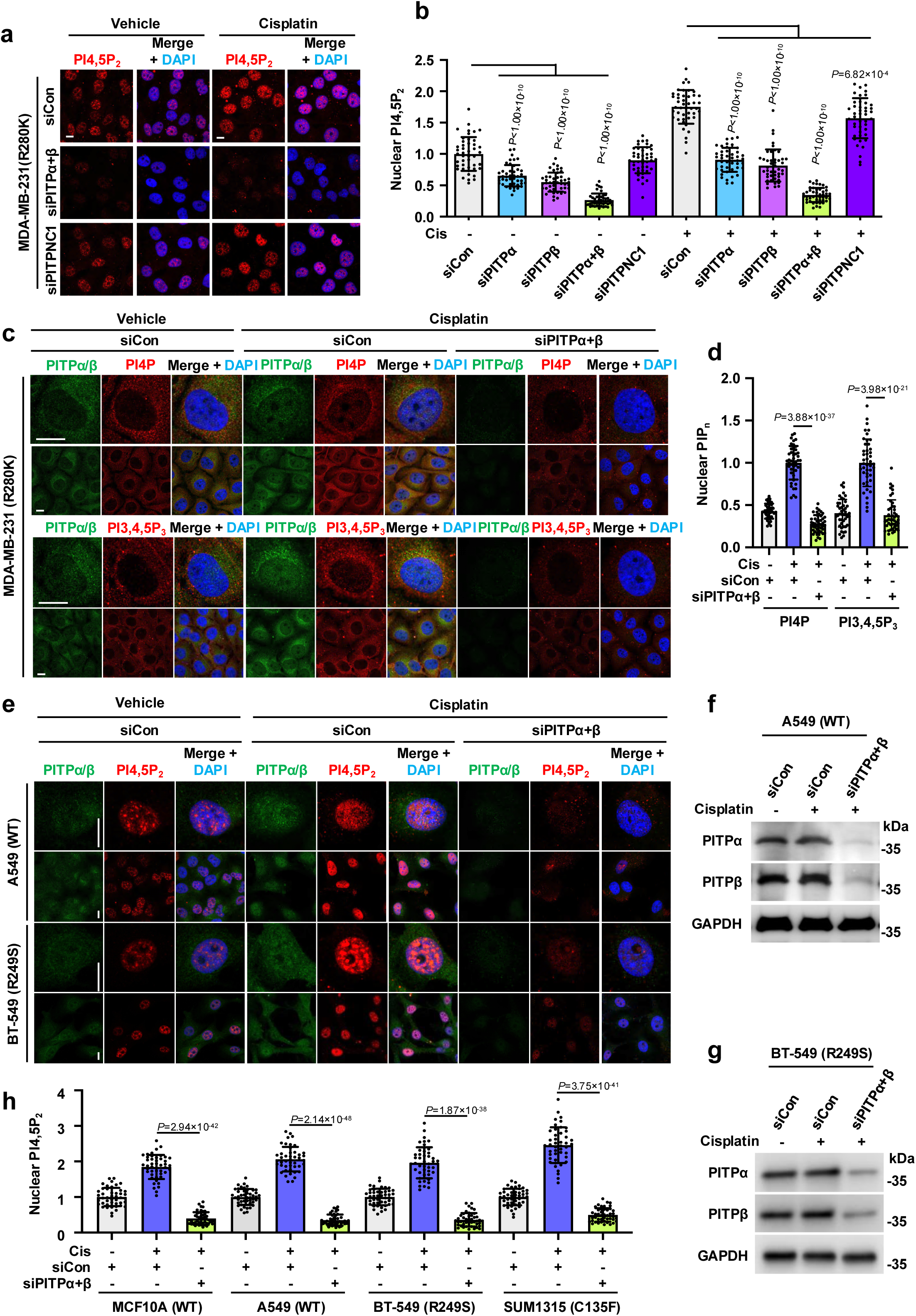
PITPα/β regulate the nuclear PIP_n_ pool. **a-b**, MDA-MB-231 cells were transfected with control siRNAs or siRNAs against PITPα, PITPβ, PITPNC1, or both PITPα and PITPβ. After 24 h, cells were treated with 30 µM cisplatin or vehicle for 24 h before being processed IF staining against PI4,5P_2_. The nuclei were counterstained by DAPI and PI4,5P_2_ levels were quantified by ImageJ (**b**). See expanded images in Extended Data Fig. 4a and KD validation in Extended Data Fig. b-c. n=3, 15 cells from each independent experiment. *p* value denotes ANOVA with Bonferroni’s multiple comparisons test. **c-d**, MDA-MB-231 cells were transfected with control siRNAs or siRNAs against both PITPα and PITPβ. After 24 h, cells were treated with 30 µM cisplatin or vehicle for 24 h before being processed IF staining against PI4P or PI3,4,5P_3_. The nuclei were counterstained by DAPI and PIP_n_ levels were quantified by ImageJ (**d**). n=3, 15 cells from each independent experiment. *p* value denotes two-sided paired t-test. **e-h**, A549 (WT), and BT-549 (R249S) cells were transfected with control siRNAs or siRNAs against both PITPα and PITPβ (**f,g**). After 24 h, cells were treated with 30 µM cisplatin or vehicle for 24 h before being processed IF staining against PI4,5P_2_. The nuclei were counterstained by DAPI and PI4,5P_2_ levels were quantified by ImageJ (**h**). n=3, 15 cells from each independent experiment. *p* value denotes two-sided paired t-test. For all graphs, data are presented as the mean ± SD. Scale bar, 5 µm

### PITPα/β control the PIP_n_ linkage to a large suite of cellular proteins

As PITPα/β regulate nuclear PIP_n_ levels and are required for p53-PIP_n_ complexes, we examined the potential role of PITPα/β in regulating potential PIP_n_ interactions with additional cellular proteins. We used PIP_n_ isomer-specific antibodies to immunoblot cell lysates by SDS-PAGE Western Blot (WB) to identify protein-PIP_n_ complexes in MDA-MB-231 breast cancer cells. Remarkably, WBs of whole cell lysates with a specific PI4,5P_2_ antibody detected many protein bands, including some that were enhanced by genotoxic stress (Fig. 3a,b), indicating that PIP_n_s stably associate with many proteins. Moreover, individual and combined targeting of PITPα/β, both KD and KO models, robustly reduced the protein-coupled PI4,5P_2_ levels in response to stress (Fig. 3a,b and Extended Data Fig. 6a). Cisplatin treatment for 24 hours achieved robust induction of protein-coupled PI4,5P_2_ levels (Extended Data Fig. 6b). Similar results were obtained in A549 lung carcinoma cells which express wild type p53 (Fig. 3c). To further support the specificity of the PI4,5P_2_ antibody, we labeled cells with [^3^H]*myo*-inositol which is specifically incorporated into PI and then phosphoinositides and inositol phosphates^33^. Consistent with the immunoblot findings, combined KD of PITPα/β also reduced [^3^H]*myo*-inositol labelling of total cellular proteins isolated by two complementary approaches, chloroform/methanol extraction followed by acetone precipitation, and analysis of proteins SDS-PAGE gels where the free PIP_n_ lipids migrate with the dye front (Fig. 3d). Furthermore, targeting of PITPα/β reduced levels of multiple protein-bound PIP_n_s (PI4P, PI4,5P_2_, PI3,4,5P_3_) and many protein bands detected by PIP_n_-specific antibodies in whole cell lysates (Fig. 3e,f), placing these PITPs as upstream regulators of PIP_n_ coupling to proteins

**Figure 3.**
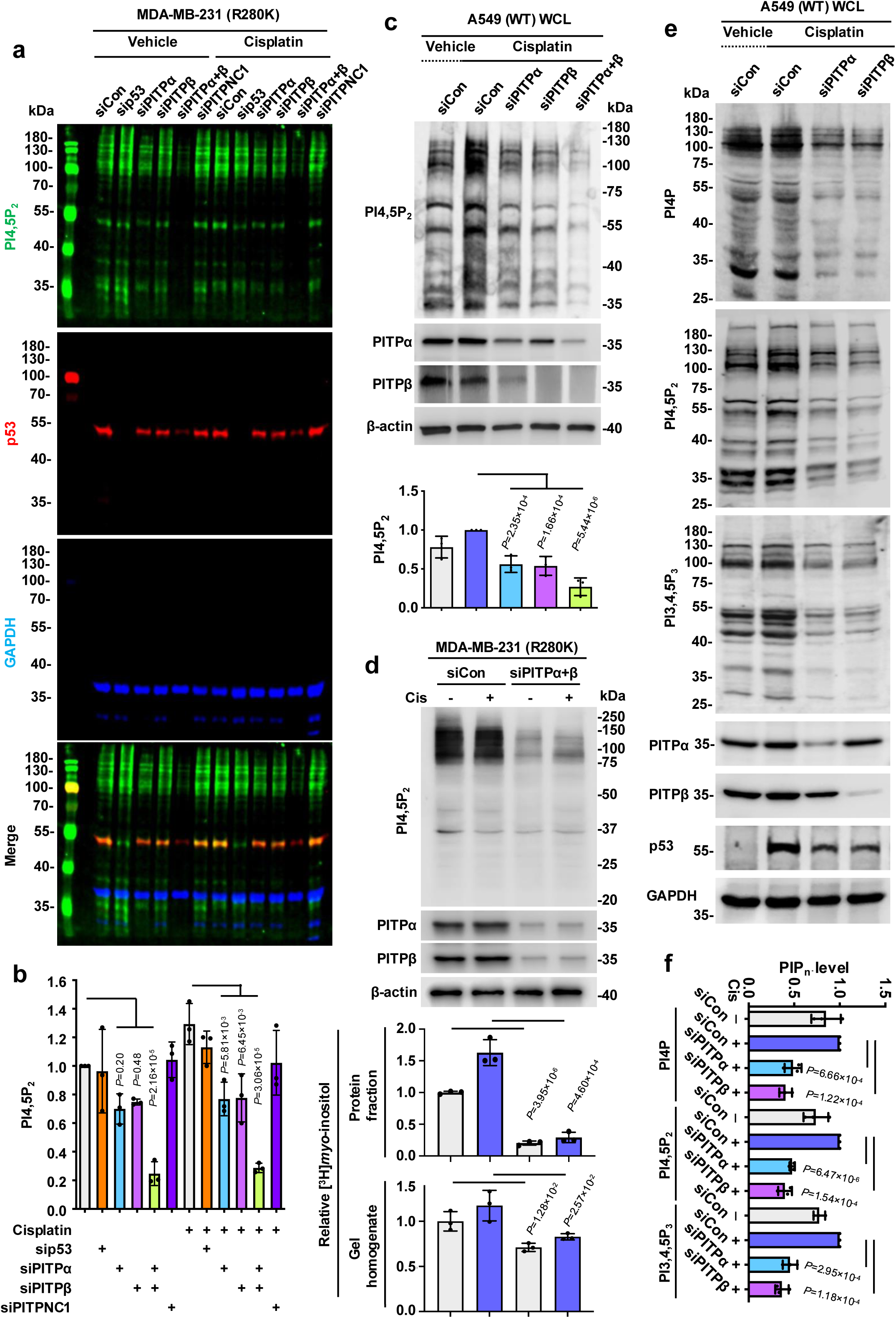
PITPα/β are required for unique and stable protein-PIP_n_ complexes. **a-b**, MDA-MB-231 cells were transfected with control siRNAs or siRNAs against p53, PITPα, PITPβ, PITPNC1, or both PITPα and PITPβ. After 24 h, cells were treated with 30 µM cisplatin or vehicle for 24 h before being processed for triple fluorescent WB for detecting p53 (red), GAPDH (blue), and protein-bound PI4,5P_2_ (green) simultaneously. The protein-bound PI4,5P_2_ levels were quantified by ImageJ (**b**). n=3 independent experiments. *p* value denotes ANOVA with Bonferroni’s multiple comparisons test. **c**, A549 cells were transfected with control siRNAs or siRNAs against PITPα, PITPβ, or both PITPα and PITPβ. After 24 h, cells were treated with 30 µM cisplatin or vehicle for 24 h before being processed for WB. The protein-bound PI4,5P_2_ levels were quantified by ImageJ. n=3 independent experiments. *p* value denotes ANOVA with Bonferroni’s multiple comparisons test. **d**, MDA-MB-231 cells were cultured from low confluency in media containing in [^3^H]*myo*-inositol or unlabeled *myo*-inositol. After 48 h, cells were transfected with control siRNAs or siRNAs against PITPα and PITPβ. 24 h later, cells were treated with 30 µM cisplatin or vehicle for another 24 h. KD and loading was confirmed by WB before the protein fraction was extracted with CHCl_3_/MeOH (top bar graph) or resolved by SDS-PAGE and the gel lane was excised (bottom bar graph). Extracted samples and dissolved gel sections were then analyzed by LSC. n=3 independent experiments. *p* value denotes two-sided paired t-test. **e-f**, A549 cells were transfected with control siRNAs or siRNAs against PITPα or PITPβ. After 24 h, cells were treated with 30 µM cisplatin or vehicle for 24 h before being processed for WB. The protein-bound PI4P, PI4,5P_2_ and PI3,4,5P_3_ levels were quantified by ImageJ (**f**). n=3 independent experiments. *p* value denotes two-sided paired t-test. For all panels, data are represented as mean ± SD.

### PI4KIIα cooperates with PITPα/β to synthesize protein-PIP_n_ complexes

As PITPα/β have been reported to cooperate with PI4Ks to synthesize PI4P^26, 34, 35^ and p53-PIP_n_ complexes (see companion paper^22^), we postulated that PI4KIIα may also regulate synthesis of other protein-PIP_n_ complexes. Indeed, KD of PI4KIIα diminished basal and stress-induced nuclear PI4,5P_2_ levels (Fig. 4a,b), similar to combined PITPα/β targeting. PI4KIIα KD also reduced protein-linked PI4,5P_2_ levels determined by WB (Fig. 4c). KD of PI4KIIα in the PITPβ KO MDA-MB-231^Cas9^ cells revealed that individual targeting of PITPα, PITPβ, or PI4KIIα is sufficient to inhibit protein-linked PIP_n_s but that combinational targeting produces more robust inhibition (Fig. 4d). Similar results were observed for the other PIP_n_s known to be protein linked: PI4P and PI3,4,5P_3_ (Fig. 4e). These finding suggest that PITP-mediated PI cargo delivery, and subsequent generation of protein-linked PI4P by PI4KIIα, are required to enable subsequent PI4,5P_2_ and PI3,4,5P_3_ linkage.

**Figure 4.**
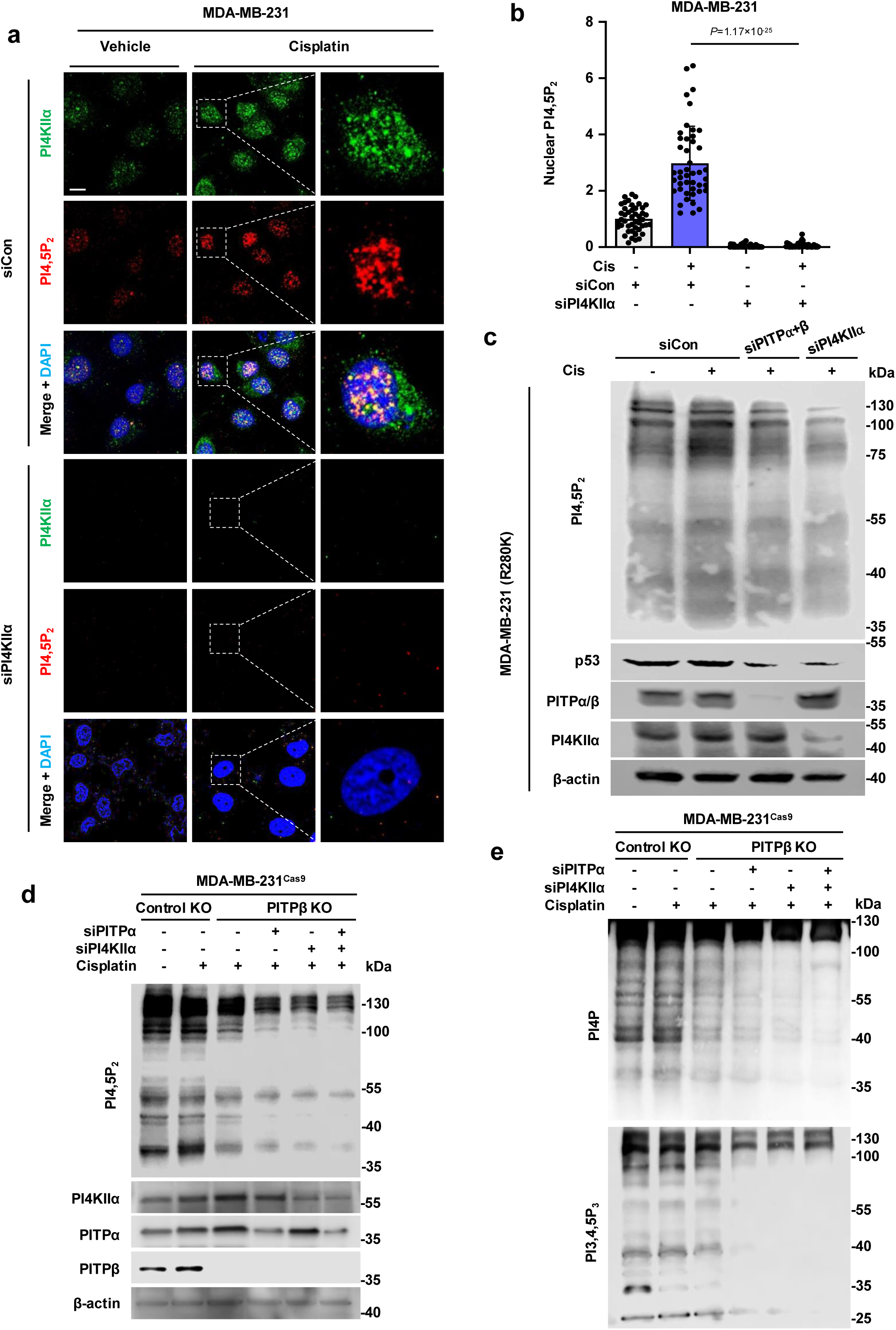
PI4KIIα is required for nuclear and protein-bound PIP_n_ signaling. **a-b**, MDA-MB-231 cells were transfected with control siRNAs or siRNAs against PI4KIIα. After 24h, cells were treated with 30 µM cisplatin or vehicle for 24 h before being processed for IF staining against PI4KIIα and PI4,5P_2_. PI4,5P_2_ levels were quantified using ImageJ (**d**). n=3, 15 cells from each independent experiment. **c**, MDA-MB-231 cells were transfected with control siRNAs or siRNAs against both PITPα and PITPβ or PI4KIIα. After 24h, cells were treated with 30 µM cisplatin or vehicle for 24 h before being processed for WB. n=3 independent experiments. **d-e**, MDA-MB-231^Cas9^ cells with PITPβ KO and control non-targeted KO were transfected with control siRNAs or siRNAs against PI4KIIα, PITPα or both PI4KIIα and PITPα. After 24 h, cells were treated with 30 µM cisplatin or vehicle for 24 h before being processed for WB against PI4,5P_2_ (**a**), PI4P (**b**), and PI3,4,5P_3_ (**b**). n=3 independent experiments. For all graphs, data are presented as the mean ± SD. Scale bar, 5 µm

### Proteomic analyses reveal potential pathways regulated by protein-PIP_n_ complexes

Given the observation that many cellular proteins form stable protein-PIP_n_ linkages, we set out to define this family of lipid-regulated proteins. First, we built on the observation that for p53, as well as Star-PAP, PIP_n_ linkages result in increased protein stability via recruitment of the small heat shock proteins αB-crystallin and HSP27^14, 20^. We successfully created double PITPα and PITPβ KO MDA-MB-231^Cas9^ cells effectively eliminating the class I PITPs from this cell line^22^. We then submitted lysates for proteomic analysis compared to a control KO in MDA-MB-231^Cas9^ cells to identify downregulated proteins in PITPα/β KO cells. Multiple downregulated proteins were identified in PITPα+β KO cells using this method, including PITPβ and p53 as internal controls to validate the approach (Fig. 5a). Many key pathways were downregulated including several major metabolic systems as well as protein export pathways (Fig. 5b). Moreover, several pathways involved in cancer metabolism and progression were also downregulated in these cells (Fig. 5b). Interestingly, glutathione and nucleotide metabolism were upregulated in the PITPα+β KO cells, perhaps as a compensatory survival mechanism (Fig. 5c).

**Figure 5.**
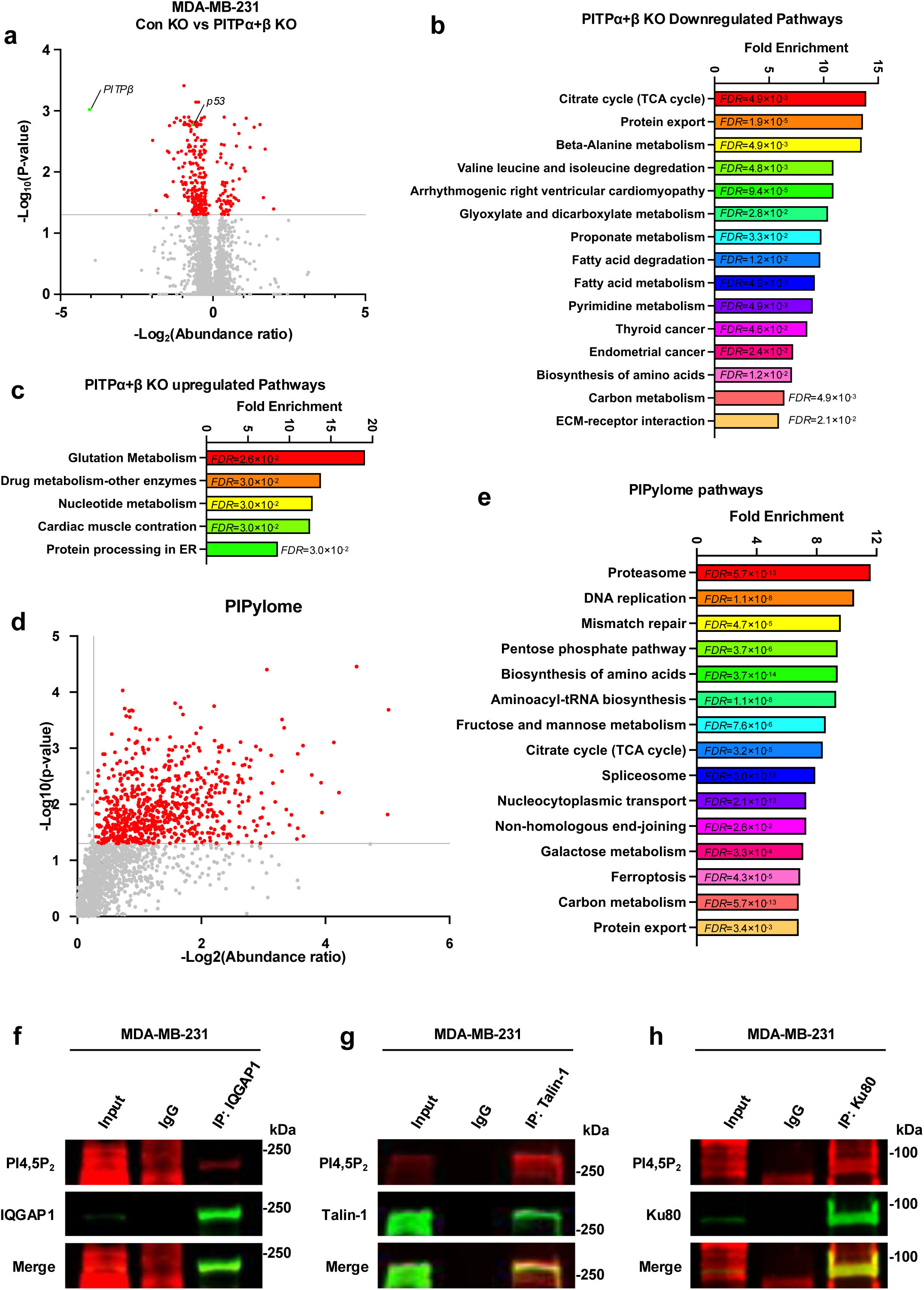
Proteomic analysis of PITPα+β KO cells and PI4,5P_2_-linked proteins. **a-c**, MDA-MB-231^Cas9^ cells harboring either a combined PITPα and PITPβ knockout or a control knockout were cultured under normal conditions. Cells were then lysed and proteins were precipitated using chloroform/methanol and resuspended before submitting for mass spectrometry proteomic analysis. All available abundance ratios and corresponding p-values were then graphed (**a**), categorized as either upregulated or downregulated, and submitted for pathway enrichment analysis using ShinyGO 0.85 (**b,c**). **d-e**, MDA-MB-231 cells were cultured under normal conditions and lysed. Then proteins were precipitated using chloroform/methanol, resuspended and incubated with anti-PI4,5P_2_ antibody. The antibody and associated proteins were then recovered using Dynabeads and submitted for mass spectrometry proteomic analysis. Identified proteins were then graphed (**d**) and submitted for pathway enrichment analysis using ShinyGO 0.85 (**e**). T-tests were used to assess significance and fold change of 0.8-1.2 were used as cut offs. **f-h**, MDA-MB-231 cells were cultured and lysed under normal conditions and IQGAP1 (**f**), Talin-1 (**g**), and Ku80 (**h**) were immunoprecipitated and resolved via WB along with PI4,5P_2_. n=3 independent experiments. For all graphs, data are presented as the mean.

Next, we developed an IP-Mass Spectrometry approach using the anti-PI4,5P_2_ antibody to identify proteins that are PI4,5P_2_-linked (“PIPylome”). Total cellular proteins were precipitated from cell lysates with chloroform/methanol to reduce competing PI4,5P_2_ lipid, dissolved in buffered SDS, diluted and combined with anti-PI4,5P_2_ antibody. The complexes were isolated by a Dyna-bead approach to enrich for PI4,5P_2_-linked proteins, eluted and analyzed by mass spectrometry (Extended Data Fig. 7a). Many proteins were identified that specifically bind the anti-PI4,5P_2_ antibody, including several that were already known to form these stable protein-PIP_n_ complexes and others that were first identified by this approach (Fig. 5d). These proteins regulate many cellular processes including DNA damage repair pathways (*e.g*., DNA replication, mismatch repair and non-homologous end-joining), metabolism (*e.g*., TCA cycle, pentose phosphate pathway, carbon and galactose), ferroptosis, signaling, regulators of cytoskeletal dynamics and motility (Fig. 5e and Extended Data Fig. 7b-e). To validate this approach, we selected three proteins that were enriched in the PIPylome. IQGAP1, Talin-1, and Ku80 all co-IP’d and retained linked PI4,5P_2_ by Western Blotting, confirming that this IP-MS approach identifies PI4,5P_2_-linked proteins (Fig. 5f-h).

## Discussion

In the companion paper, we discovered that the lipid transfer proteins PITPα/β bind wild-type and mutant p53, and together with the PIP kinase PI4KIIα, initiate the synthesis of p53-PIP_n_ complexes^22^. The linked PIP_n_ isomers regulate p53 stability and function^13, 14^. Moreover, PIP_n_s are known to stably associate with several other proteins, including Star-PAP, NRF2 and MDM2^19, 21, 36^. This prompted us to investigate whether PITPs and PI4KIIα play a broader role in linking PIP_n_s to other proteins. Remarkably, WB of whole cell lysates from multiple cell lines with antibodies specific for PIP_n_ isomers revealed many proteins with stably associated PIP_n_s, fundamentally distinct from conventional PIP_n_ protein binding. Notably, metabolic labeling of cells with the phosphoinositide precursor [H^3^]-*myo*-inositol resulted in the stable incorporation of [H^3^] label into the protein bands despite harsh denaturing conditions, further underscoring that the PIP_n_s are protein linked. Furthermore, both the protein bands identified by WB with PIP_n_ antibodies and by [H^3^]-*myo*-inositol incorporation are reduced by individual KD/KO of PITPα/β, while combined KD/KO of PITPα/β resulted in more robust suppression. Importantly, PI binding by PITPα is required to rescue the effects of PITPα/β KD/KO, highlighting that this well-described function of these transfer proteins is required to initiate PIP_n_ linkage to proteins. PI4KIIα KD mimics the effects of PITPα/β KD/KO, indicating dual regulation of these PIP_n_-linked proteins. Consistent with this newly discovered function of PITPs, their cellular localization responds dynamically to diverse cellular stressors, resulting in their accumulation in the nucleus where they generate a PIP_n_ pool, which is likely anchored to proteins. Overall, these data support the existence of a unique panel of proteins that are PIP_n_-linked, the “PIPylome”, by a mechanism that requires the concerted action of PITPα/β and PI4KIIα. We are actively investigating the chemical nature and amino acid(s) localization of this linkage in p53 and other modified proteins.

As a first step in delineating the PITPα/β-regulated PIPylome, we utilized two orthogonal approaches: 1) an analysis of the proteome in control versus combined PITPα/β KO cells; and 2) an analysis of PI,4,5P_2_-associated proteins. The first approach stems from our observation that PIP_n_-linkage regulates the stability of known members of the PIPylome including p53, MDM2, Star-PAP, YAP/TAZ and NRF2 ^13, 14, 16, 19, 21, 36^. Pathway analysis identified many protein components of metabolic pathways that are differentially expressed in the presence and absence of PITPα/β, a subset of which are also concordantly enhanced in the PI4,5P_2_ interactome (*e.g*., TCA cycle, carbon metabolism, and protein export). We validated a small number of PI4,5P_2_ interacting proteins by IP and subsequent WB with PI4,5P_2_ antibody. IQGAP1 is a cytoplasmic protein that scaffolds the membrane-localized PI3K/Akt pathway and regulates cell migration^6, 37^, suggesting the possibility that PIP_n_ linkage to IQGAP1 may regulate the assembly and/or activation of canonical PI3K/Akt signaling and/or cell migration. Talin interacts with the PI4,5P_2_-generating lipid kinase PIPKIγ to regulate cell adhesion, invasion and epithelial-mesenchymal transition^38–40^, suggesting a role of PIP_n_-linked Talin in cell invasion, migration and metastasis. Additionally, the observation that both the α and γ isoforms of PIPKI bind and phosphorylate PIP_n_-linked proteins points to the likely versatility of this pathway. Ku80 and Ku70 form a heterodimer to recruit DNA-PK to sites of DNA damage^41, 42^, suggesting a functional role of PIPylation in the DNA damage response. Indeed, DNA damage induces the rapid accumulation of PIP_n_s in the nucleus and have been proposed to regulate ATR signaling^43, 44^ and p53^13, 14^ in the DNA damage response^43^. The observation that multiple DNA damage pathway components are in the PI4,5P_2_ interactome strongly suggests a functional role for PIPylation in this process

In summary, we have provided evidence that the PIP_n_ isomers (PI4P, PI4,5P_2_ and PI3,4,5P_3_) are stably linked to proteins in the nucleus and cytoplasm. Given the existence of seven distinct PIP_n_ isomers^4, 5^, it seems plausible that these other PIP_n_ isomers are also linked to proteins and modulate signaling. Notably, these protein-linked PIP_n_s are signaling competent as they are modified by specific PIP kinases and phosphatases and each isomer selectively recruits a set of downstream effectors, *e.g*., small heat shock proteins, Akt-activating kinases and Akt substrates^4, 5, 13, 14, 19, 21, 36^. Significantly, the interactions of p53 with these PIP metabolizing enzymes and Akt pathway components occurs in the absence of the PIP_n_ linkage (with *E. coli* expressed p53)^13, 14^. This indicates that in the absence of PIP_n_ linkages, protein-protein interactions provide the functional specificity for p53 and likely other proteins^16, 19, 21, 36^. The linkage of PIP_n_ second messengers to proteins represent a “third messenger” pathway that functions independently of the canonical membrane-localized pathway and fine tunes the interactions and functions of the linked protein, the details of which are just beginning to emerge.

## ACKNOWLEDGMENTS

We thank Dr. Adrea Galmozzi for discussions and comments, and Lance Rodenkirch for technical support. M.C. is currently supported by grant D2301007 from the Shenzhen Medical Research Fund. This work was supported in part by a National Institutes of Health grant R35GM134955 (R.A.A.), R01CA286492 (R.A.A. & V.L.C.), Department of Defense Breast Cancer Research Program grants W81XWH-17-1-0258 (R.A.A.), W81XWH-17-1-0259 (V.L.C.), W81XWH-21-1-0129 (V.L.C.), HT9425-23-1-0553 (V.L.C.), and HT9425-23-1-0554 (R.A.A.), and a grant from the Breast Cancer Research Foundation (V.L.C.).

## AUTHOR CONTRIBUTIONS

N.D.C, M.C., P.A., T.W., B.B.M., M.S., V.L.C., and R.A.A. designed the experiments and project. N.D.C., M.C., P.A., T.W., W.C., G.V., B.B.M., and D.B. performed the experiments. N.D.C., P.A., V.L.C., and R.A.A. wrote the manuscript. M.C., W.C., G.V., B.B.M., and M.S. reviewed and revised the manuscript.

## Experimental Procedures

### Cell culture and constructs

All cell lines including A549, BT-549, Cal33, HCT116, HS578T, MDA-MB-231, SUM159, SUM1315, HEK293FT, and MCF-10A cells were purchased from ATCC. MCF10A cells were grown in DMEM/F12 (#11330-032, Invitrogen) with supplements of 10% fetal bovine serum (#100-106, GeminiBio), 1% penicillin/streptomycin (#15140-122, Gibco), 20 ng/mL EGF (#CC-4107, Lonza), 0.5 mg/mL Hydrocortisone (#H4001, Sigma), 100 ng/mL Cholera toxin (#C-8052, Sigma), and 10 μg/mL Insulin (#I-1882, Sigma). The remaining cell lines were maintained in DMEM (#10-013-CV, Corning) supplemented with 10% fetal bovine serum (#100-106, GeminiBio) and 1% penicillin/streptomycin (#15140-122, Gibco). The cell lines used in this study were routinely tested for mycoplasma using MycoAlert Kit (#LT07-318, Lonza), and mycoplasma-negative cells were used. None of the cell lines used in this study are listed in the database of commonly misidentified cell lines maintained by ICLAC. The HA-tagged wild-type PITPα construct and PI-binding deficient mutant T59D^45^ were purchased from Genscript. Plasmids were transfected according to the manufacturer’s instructions into mammalian cells using Lipofectamine^TM^3000, (#L3000015, Thermo Fisher Scientific). Typically, 2-5 µg of DNA was used alongside 6-10 µl of lipid in 6-well plates for transfection. Cells were selected for at least 80% transfection efficiency and used for further analysis.

### Antibodies and reagents

The monoclonal antibodies that were used are against p53 (clone DO-1, #SC-126, Santa Cruz Biotechnology), p53 (clone 7F5, #2527, Cell Signaling), pAkt^S473^ (clone 193H12, #4058, Cell Signaling), HA-tag (clone C29F4, #3724, Cell Signaling), GAPDH (clone 0411, #sc-47724, Santa Cruz Biotechnology), β-actin (clone 13E5, #4970, Cell Signaling), Lamin B2 (clone D8P3U, #12255, Cell Signaling), IQGAP1 (clone D-3, #sc-374307, Santa Cruz Biotechnology), Talin-1 (clone C-9, #sc-365875, Santa Cruz Biotechnology), Ku80 (clone C48E7, #2180, Cell Signaling), and polyclonal antibodies against PITPα (#16613-1-AP, ThermoFisher), PITPβ (#ab127563, abcam), PITPNC1 (IF, #ab222078, abcam), PITPNC1 (WB, #NBP2-19842, Novus), PI4KIIα (#NBP2-44158, Novus). Anti-PI4P (#Z-P004), PI4,5P_2_ (#Z-P045) and PI3,4,5P_3_ (#Z-P345) antibodies were used for immunostaining, proximity ligation assay (PLA), and WB analyses. For immunoblotting analyses, all antibodies were diluted at a 1:1000 ratio except for GAPDH (clone 0411, 1:5000) and p53 p53 (clone DO-1, 1:5000). For protein immunoprecipitation, antibody-conjugated agarose was purchased from Santa Cruz Biotechnology, including agarose-conjugated antibodies against IQGAP1 (#sc-374307AC), Talin-1 (#sc-365875AC), and Ku80 (#sc-5280AC). For immunostaining analyses and proximity ligation assay PLA, all primary antibodies were diluted at a 1:100 ratio. Nuclear envelope marker (Alexa Fluor**®**488 Lamin A/C, clone 4C11, #8617, Cell Signaling, 1:200) was used to identify the nuclear boundry. For the knockdown (KD) experiments, the ON-TARGETplus siRNA SMARTpool with 4 siRNAs in combination against human PITPα (#L-018010-00), PITPβ (#L-006459-01), PITPNC1 (#L-012476-02), and PI4KIIα (#LQ-006770-00-0020) were purchased from Dharmacon. Non-targeting siRNA (#D-001810-01, Dharmacon) was used as a control. For the KD and rescue experiments, methodology was the same as in the companion manuscript for this publication^22^ and KD efficiency was determined by immunoblotting. KD efficiency greater than 80% was required to observe phenotypic changes in the study. Cisplatin, (#NC1706394, Fisher Scientific), was used as cellular stressors.

### Immunoprecipitation and immunoblotting

Immunoprecipitation methodology was performed the same as found and described in the companion manuscript for this publication^22^. Immunoblots were developed by Odyssey Imaging System (LI-COR Biosciences) and protein levels were quantified using ImageJ. The unsaturated exposure of immunoblot images was used for quantification with the appropriate loading controls as standards. Statistical data analysis consisted of T-test and ANOVA and was performed with Microsoft Excel and PRISM, respectivley, using data from at least three independent experiments.

### Fluorescent IP-WB

Cells were lysed in a RIPA lysis buffer system^13^ after the indicated treatment and quantified for protein concentration as described above. For endogenous protein immunoprecipitation, 0.5-1 mg of cell lysates were incubated with 20 µl anti-IQGAP1 (clone D-3, #sc-374307AC, Santa Cruz Biotechnology), Talin-1 (clone C-9, #sc-365875AC, Santa Cruz Biotechnology), or Ku80 (clone B-1, #sc-5280AC, Santa Cruz Biotechnology) monoclonal IgG antibody-conjugated agarose at 4°C for 24 h. Normal immunoglobulin (IgG)-conjugated agarose was used as a negative control (#sc-2343, Santa Cruz Biotechnology). IP methodology is the same as described in the companion manuscript for this publication^22^. For double fluorescent IP-WB detecting protein-PI4,5P_2_ complexes, anti-IQGAP1 (#sc-374307), -Talin-1 (#sc-365875), or -Ku80 (#2180) rabbit monoclonal IgG antibody at 1:2000 dilution and anti-PI4,5P_2_ mouse monoclonal IgM antibody (#Z-P045, Echelon Biosciences) at 1:1000 dilution were mixed together in blocking buffer with 0.02 % Sodium Azide and incubated with the membrane at 4°C overnight. The next day, the membrane was thrice washed with TBST for 10 min. Secondary antibody incubation and imaging were conducted as described in the companion manuscript^22^. Statistical data analysis was performed with Microsoft Excel, using data from at least three independent experiments.

### Immunofluorescence (IF) and Confocal Microscopy

For immunofluorescence studies, methodology, statistical testing, quantification and correlation analysis was conducted as described in the companion manuscript^22^. A value of 1 represents perfect correlation, 0 means no correlation, and −1 means perfect negative correlation^46^. Pearson’s r greater than 0.7 suggests a strong correlation^47^.

### [^3^H]*myo*-inositol Metabolic Labeling

Radio-labeling experiments were conducted as described in the companion manuscript for this publication^22^.

### Lenti-CRISPR Gene Editing

All gene editing was conducted as described in the companion manuscript for this publication^22^.

### Mass Spectrometry proteomics of MDA-MB-231 PITPα+β KO cells

MDA-MB-231 PITPα+β KO cells were generated as described above in *Lenti-CRISPR Gene Editing*. Parental and KO cells were plated in triplicate and grown to ∼80% confluency after 24 h. The plates were then lysed using RIPA and batched together to minimize variability before MeOH:CHCl_3_ (2:1) was added to form protein precipitate. Soluble phases in MeOH and CHCl_3_ were aspirated out and the protein flake was dried using a speed vacuum. Pellets were resolubilized into 500µL 8M urea (#U15, Fisher) in 50mM ammonium bicarbonate (#A643, Fisher), pH 8.5. 25µL of this was taken and diluted to 4M urea with 50mM ammonium bicarbonate. Samples were reduced with 2mM TCEP (#C4706, Sigma) for 45 minutes at 42 C, then alkylated with 5mM iodoacetamide (#I1149, Sigma) for 45 minutes in the dark at room temperature, then quenched with another aliquot of 2mM TCEP for 5 minutes. Samples were diluted to 1M urea with 50mM ammonium bicarbonate and proteolytically digested for 12 hours at 37°C using a 1:1 mixture of Trypsin:Lys-C (#V5071, Promega) and a protease:protein ratio of 1:50. Samples were acidified by addition of neat formic acid (#A117, Fisher) to 1% vol/vol, desalted and concentrated using OMIX C18 tips (#A57003100, Agilent) and manufacturer’s protocol, and dried to completion with a vacuum centrifuge. Resulting dried down peptides were resolubilized into 50µL 0.1% formic acid (#LS118, Fisher) and 3µM of this was injected for spectral analysis.

The liquid chromatography (LC) system used was a Dionex UltiMate 3000 and the LC conditions were as follows: buffer A is 0.1% formic acid (#LS118, Fisher), buffer B is 80/20/0.1% acetonitrile/water/formic acid, flow rate was 300nL/min, and the column used was a 50cm Thermo Scientific PepMap RSLC C18 column with 2µm bead size, 100Å pore size, and 75µm inner diameter. After column loading in 2% B, peptide samples were eluted with a 75-minute linear gradient from 0-25%B, followed by a 20-minute linear gradient from 25-50%B, a 4 min linear gradient to 95%B, flushing with 05% B for 3 min, then requilibration to 2% B for 15 minutes. MS1 acquisition conditions were as follows: cycle time of 1s between MS1 scans, acquisition in the orbitrap mass analyzer with a resolution of 120K, scan range of 350-1600m/z, normalized AGC target of 250%, profile data and positive mode, and a max inject time of 50ms. Fragment ion selection was filtered using MIPS mode set to peptide and charge state set to 2-6. Dynamic exclusion was used with an n of 1 for 10 seconds, and a mass tolerance +/− 10ppm. MS2 spectra were acquired in the ion trap with quadrupole isolation of 0.7, HCD activation/fragmentation with 30% collision energy, turbo scan rate, max inject time of 25ms, normalized AGC target of 300%, scan range set to auto, and positive mode with centroid data. Raw data was searched with the software Proteome Discoverer v2.4. The H. Sapiens Uniprot proteome (20,353 sequences) was searched using the Sequest algorithm alongside a database of common contaminants with the following parameters: a precursor mass tolerance of 10ppm, fragment ion mass tolerance of 0.6Da, max missed cleavages of 2, carbamidomethylation (+57.02) set as a fixed modification on cysteines. The following modifications were set as dynamic: oxidation (+15.99) on methionines, deamidation (+0.98) on asparagines and glutamines, and phosphorylation on serines, threonines, and tyrosines (+79.97). A concatenated database search strategy was used in the Percolator node, and an FDR of 0.05 was set. For scoring and further filtering, all default parameters were used. For label-free quantification in Proteome Discoverer, peptide precursor abundances based on intensity and summed abundances were used. Modified peptides were not included in quantification.

### Mass Spectrometry proteomics of anti-PI4,5P_2_ immunoprecipitation

MDA-MB-231 cells were plated in triplicate and allowed to reach ∼80% confluency. The cells were then lysed using a RIPA lysis system (#sc-24948, Santa Cruz Biotechnology) with 1 mM Na3VO4, 5 mM NaF, and 1x protease inhibitor cocktail (#11836153001, Roche). The triplicate groups were then compiled into a single batched lysate and proteins were precipitated using a 4:1 ratio of Chloroform:Methanol. The protein precipitate was resuspended in 3% SDS in PBS and split into 6 tubes; 3 tubes were incubated with 10 μg anti-PI4,5P2 IgM antibody (#Z-P045, Echelon Biosciences) and 3 tubes were incubated with normal IgM antibody (sc-3881, Santa Cruz) for 24 h while rotating. After 24 h, the resuspended proteins were added to pre-cleared anti-mouse IgM Dynabeads (#11039D, Invitrogen) and rotated at room temperature for 1.5 h. The bound complexes were then purified via magnet and bound proteins were eluted using 0.1 M citrate. Eluates were brought to 5% final SDS. Samples were reduced for 35 min with 2mM Tris(2-carboxyethyl)phosphine hydrochloride (TCEP, Sigma, #C4706) at 42°C, alkylated for 35 min with iodoacetamide (#I1149, Sigma) at room temperature in the dark, then quenched with a further 2mM TCEP for 5 min at room temperature. Samples were digested on S-Trap Micro digestion columns (Protifi) per manufacturer’s protocol with 3µg of Trypsin/Lys-C mix (#V5071, Promega). Resulting dried down peptides were resolubilized into 10µL 0.1% formic acid (#LS118, Fisher) and 1.5µL of this was injected for spectral analysis as described above in *Mass Spectrometry proteomics of MDA-MB-231 PITPα+β KO cells*.

### Statistics and Reproducibility

One-way ANOVA was used for group significance with Bonferroni’s multiple comparisons test used for significance within grouped samples and Two-tailed unpaired *t*-tests were used for pair-wise significance. In this study, no power calculations were used. Experimental design and sample sizes were determined based on previously published experiments where significance was readily observed^13, 14^. As noted, each experiment was independently repeated at least three times, and the number of repeats is defined in the figure legend. We used at least three independent experiments or biologically independent samples for statistical analysis.

### Resource and Data Availability

All data supporting the findings of this study are available from the corresponding authors on reasonable request.

---

**Extended Data Figure 1.**
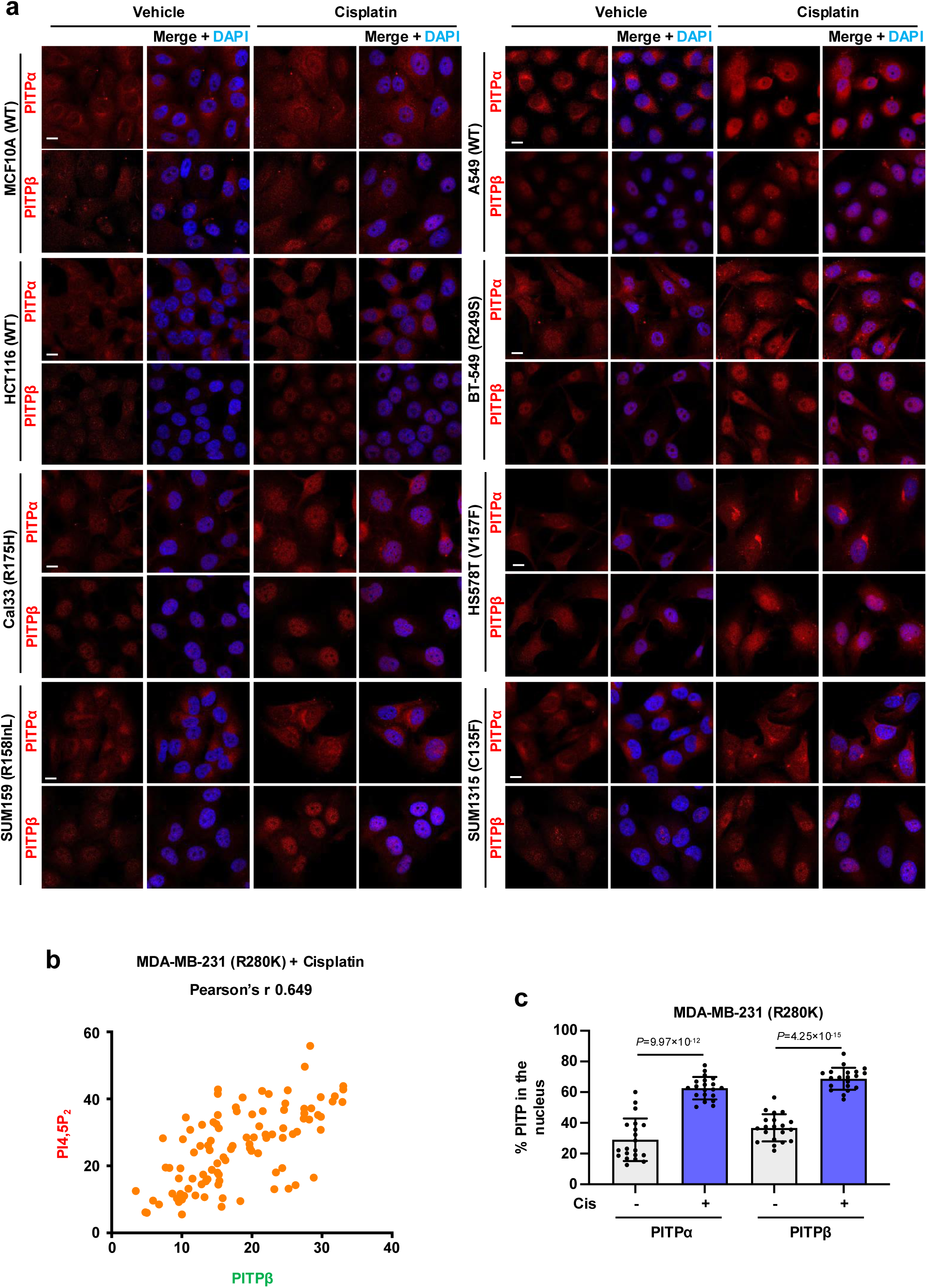
PITPα/β nuclear accumulation in multiple cell lines. **a**, Confocal IF of PITPα or PITPβ in MCF10A (WT), A549 (WT), HCT116 (WT), BT-549 (R249S), Cal33 (R175H), HS578T (V157F), SUM159 (R158InL), and SUM1315 (C135F) cells treated with vehicle or 30 µM cisplatin for 24 h. The nuclei were counterstained by DAPI. See quantification in Fig. 1c. n=3 independent experiments. **b**, Correlation analysis using LASX between PITPβ and PI4,5P_2_ levels as determined by IF in MDA-MB-231 cells treated with 30 µM cisplatin for 24 h. Pearson’s r=0.649. **c**, Localization analysis of PITPα/β as determined by IF in MDA-MB-231 cells treated with 30 µM cisplatin for 24 h. Nuclear regions were defined by DAPI counterstaining and nuclear IF signals were divided by total IF signals for individual cells. n=3 independent experiments. For all graphs, data are presented as the mean ± SD. Scale bar, 5 µm.

**Extended Data Figure 2.**
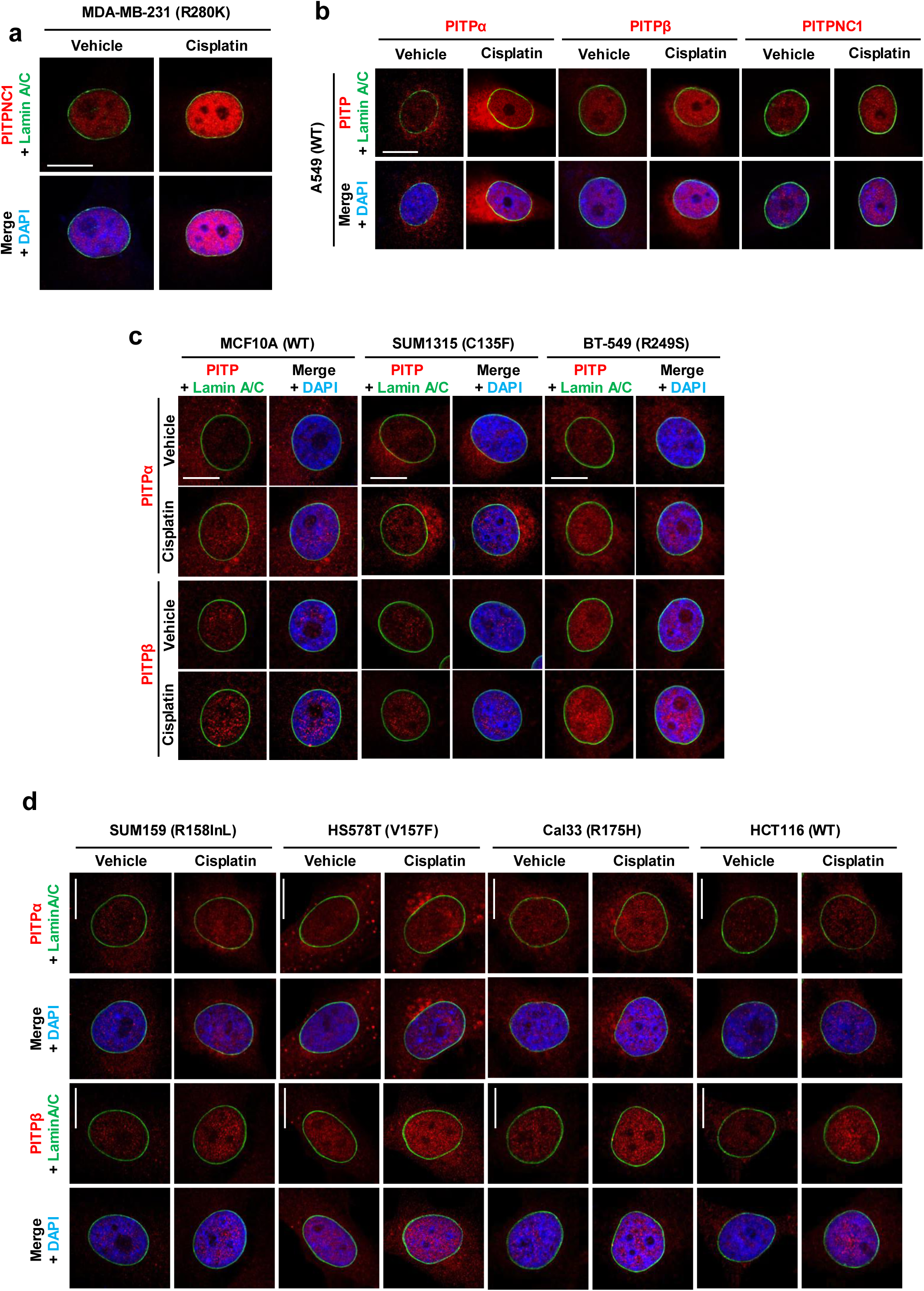
PITPα/β accumulate in the non-membranous nucleoplasm. **a**, Confocal IF of PITPNC1 overlaid with the nuclear envelope marker Lamin A/C in MDA-MB-231 cells treated with vehicle or 30 µM cisplatin for 24 h. The nuclei were counterstained by DAPI. See PITPα and PITPβ IF in Fig. 1d. n=3 independent experiments. **b**, Confocal IF of PITPα, PITPβ, or PITPNC1 overlaid with the nuclear envelope marker Lamin A/C in A549 cells treated with vehicle or 30 µM cisplatin for 24 h. The nuclei were counterstained by DAPI. n=3 independent experiments. **c**, Confocal IF of PITPα or PITPβ overlaid with the nuclear envelope marker Lamin A/C in MCF10A (WT), SUM1315 (C135F), and BT-529 (R249S) cells treated with vehicle or 30 µM cisplatin for 24 h. The nuclei were counterstained by DAPI. n=3 independent experiments. **d**, Confocal IF of PITPα or PITPβ overlaid with the nuclear envelope marker Lamin A/C in SUM159 (R158InL), HS578T (V157F), Cal33 (R175H), and HCT116 (WT) cells treated with vehicle or 30 µM cisplatin for 24 h. The nuclei were counterstained by DAPI. n=3 independent experiments. Scale bar, 5 µm.

**Extended Data Figure 3.**
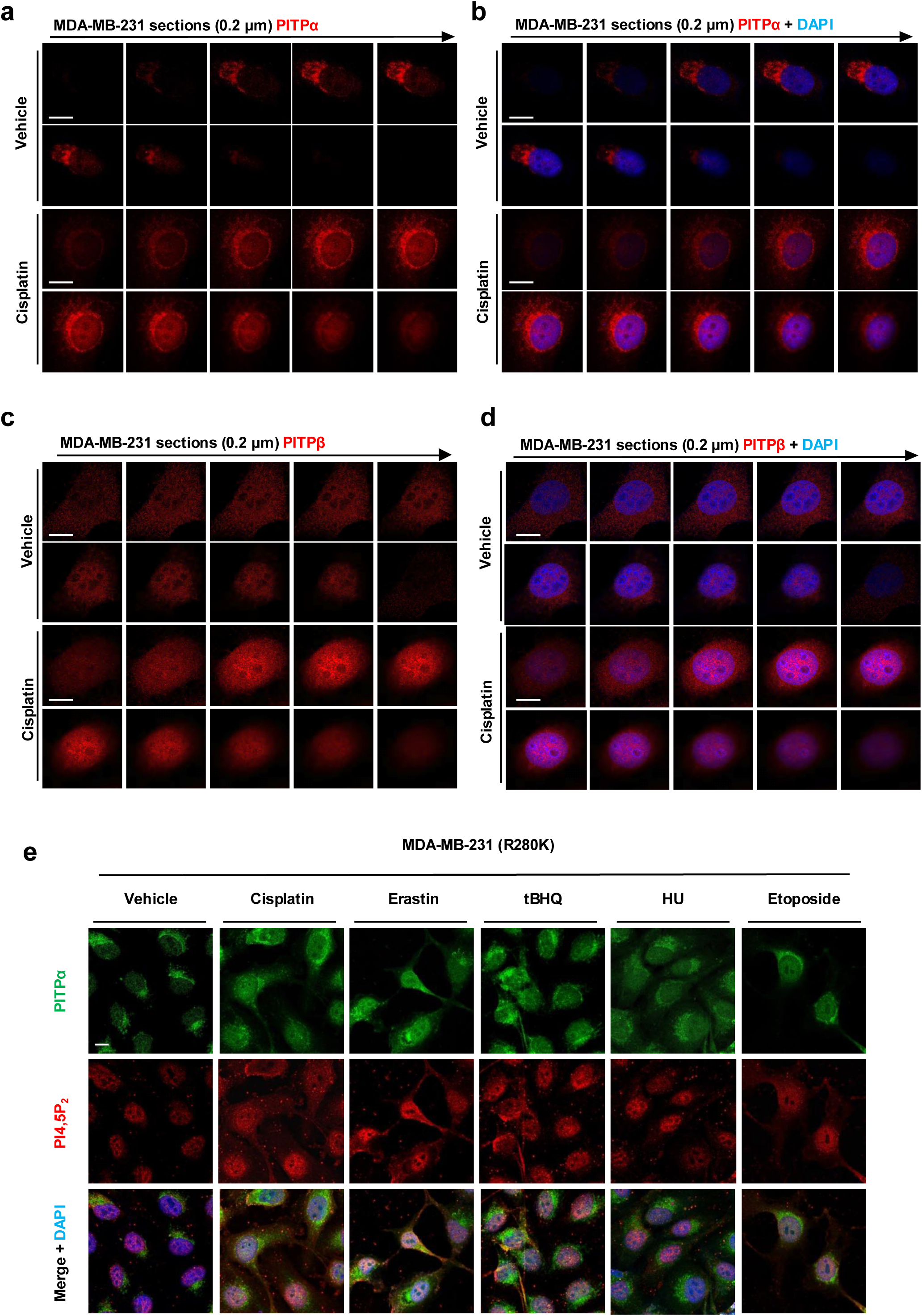
3D sectioning of PITPα/β nuclear accumulation. **a-b**, 3D sections of IF staining against PITPα in MDA-MB-231 cells treated with vehicle or 30 µM cisplatin for 24 h. The nuclei were counterstained by DAPI. Each frame of the 3D sections was over a 0.2 μm thickness. **c-d**, 3D sections of IF staining against PITPβ in MDA-MB-231 cells treated with vehicle or 30 µM cisplatin for 24 h. The nuclei were counterstained by DAPI. Each frame of the 3D sections was over a 0.2 μm thickness. **e**, MDA-MB-231 cells treated with 30 µM cisplatin, 20 µM erastin, 100 µM tBHQ, 100 µM etoposide, 100 µM hydroxyurea, or vehicle for 24 h. The cells were processed for IF staining against PITPα and PI4,5P2. The nuclei were counterstained by DAPI. The nuclear levels of PITPα and PI4,5P2 were quantified by ImageJ (**f**). See quantification in Fig. 1f. n=3, 15 cells from each independent experiment. Scale bar, 5 µm.

**Extended Data Figure 4.**
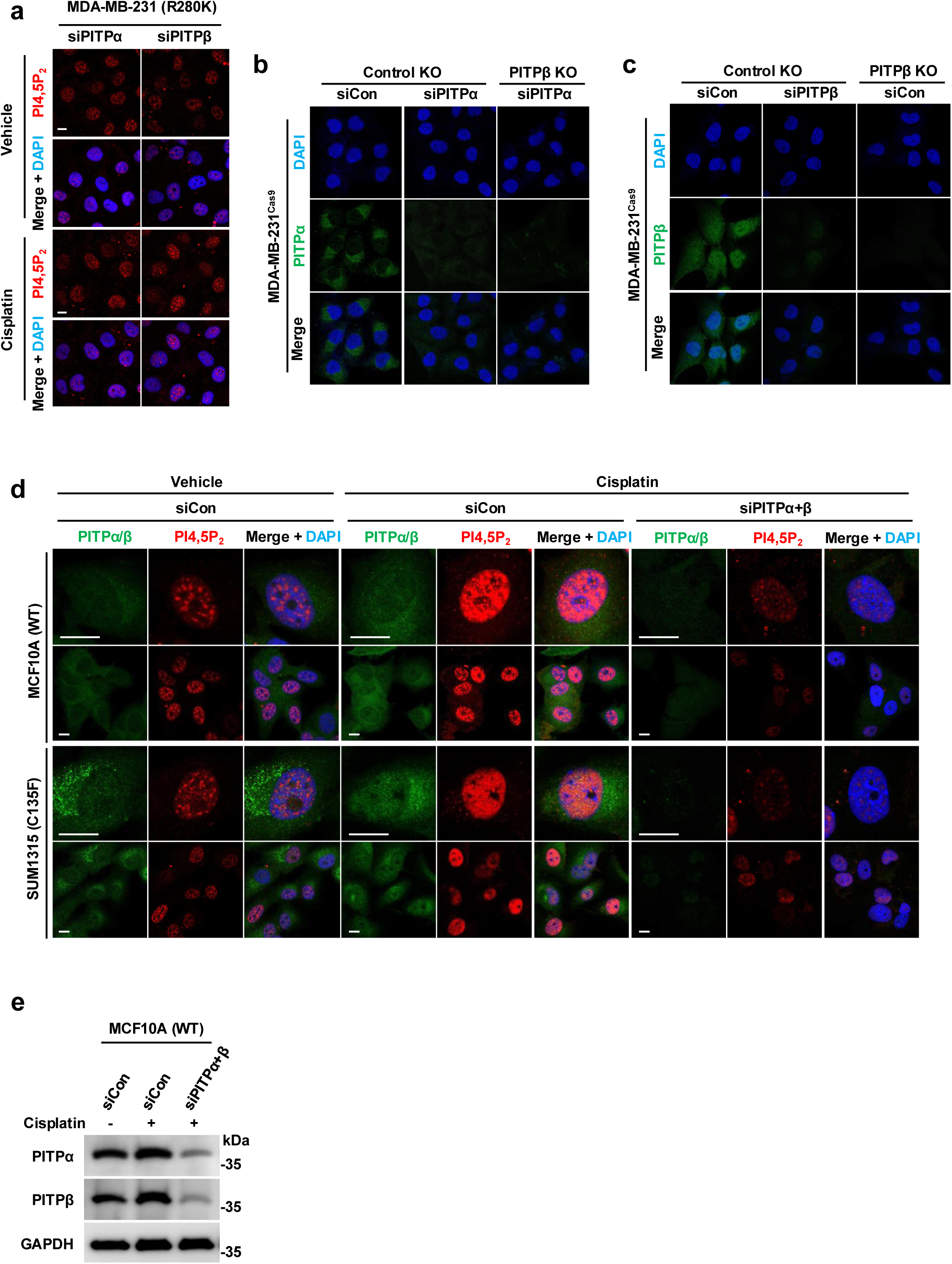
PITPα/β are required for nuclear PI4,5P_2_ pools. **a**, MDA-MB-231 cells were transfected with siRNAs against PITPα or PITPβ. After 24 h, cells were treated with 30 µM cisplatin or vehicle for 24 h before being processed IF staining against PI4,5P_2_. The nuclei were counterstained by DAPI. See expanded images in Fig. 2a and quantification in Fig. 2b. n=3 independent experiments. **b-c**, MDA-MB-231^Cas9^ cells harboring either a PITPα knockout, PITPβ knockout, or a control knockout were transfected with control siRNA or siRNAs against PITPα or PITPβ. After 48 h cells were processed for IF staining against PITPα or PITPβ. n=3 independent experiments. **d**, MCF10A (WT) and SUM1315 (C135F) cells were transfected with control siRNAs or siRNAs against both PITPα and PITPβ. After 24 h, cells were treated with 30 µM cisplatin or vehicle for 24 h before being processed IF staining against PI4,5P_2_. The nuclei were counterstained by DAPI. See quantification in Fig. 2h. n=3 independent experiments. Scale bar, 5 µm.

**Extended Data Figure 5.**
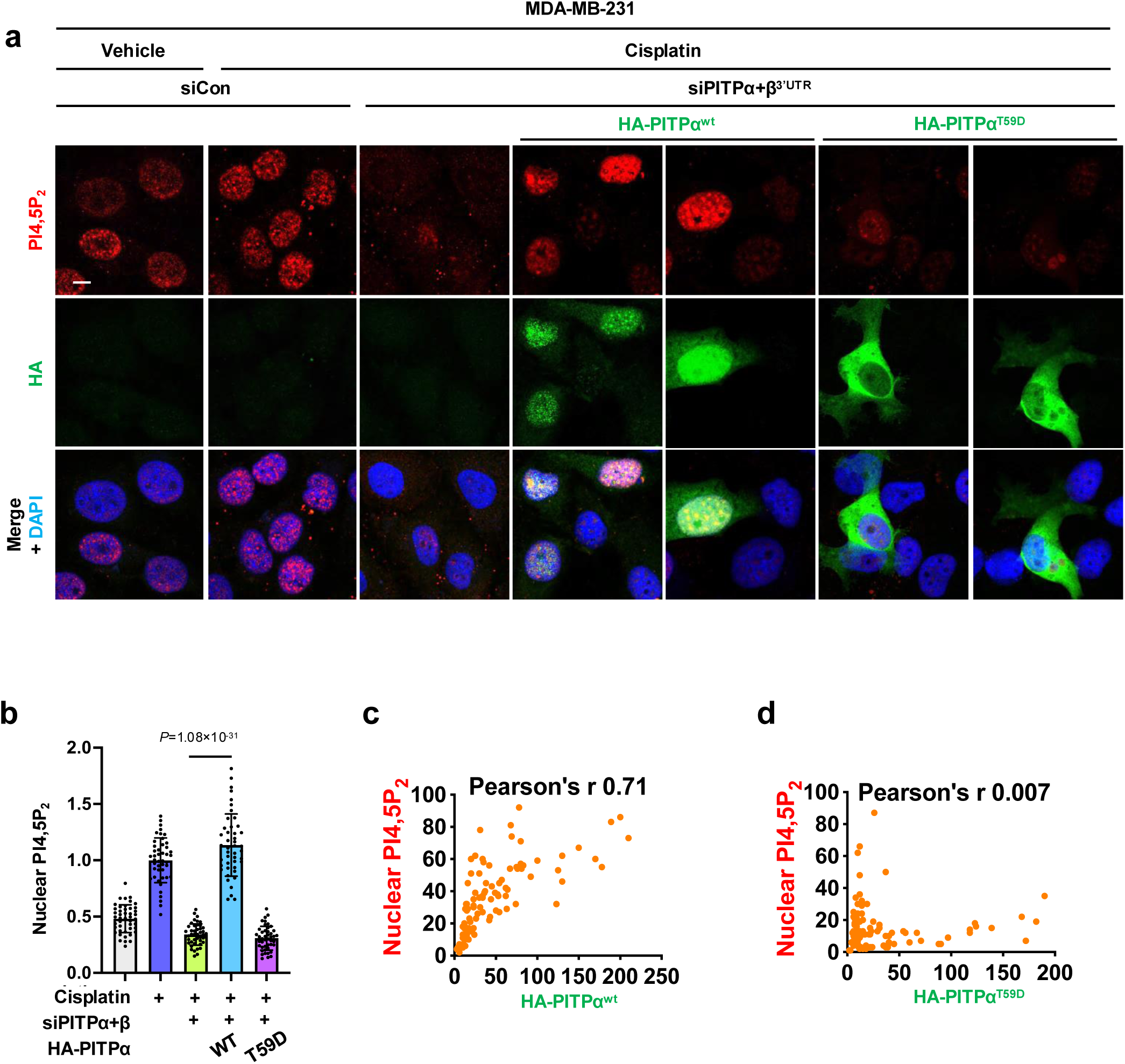
PI binding is required for PITP-dependent nuclear PI4,5P_2_. **a-b**, MDA-MB-231 cells were transfected with control siRNAs or siRNA against the 3’UTR of both PITPα and PITPβ for 24 h. Then the cells were transfected with either HA-tagged wild-type PITPα or PI-binding defective T59D PITPα. After 24 h, cells were treated with 30 μM cisplatin of vehicle for 24 h before being processed for IF against HA-tag and PI4,5P_2_ (**a**). The nuclear PI4,5P_2_ levels in HA-positive cells were quantified by ImageJ (**b**). n=3 independent experiments. **c-d**, Correlation analysis using LASX between HA-tag (PITPα^wt^ (**c**) or PITPα^T59D^ (**d**)) and PI4,5P_2_ levels as determined by IF in MDA-MB-231 cells treated with 30 µM cisplatin for 24 h. Pearson’s r=0.71 and 0.007 for PITPα^wt^ and PITPα^T59D^, respectively. For all graphs, data are presented as the mean ± SD. Scale bar, 5 µm.

**Extended Data Figure 6.**
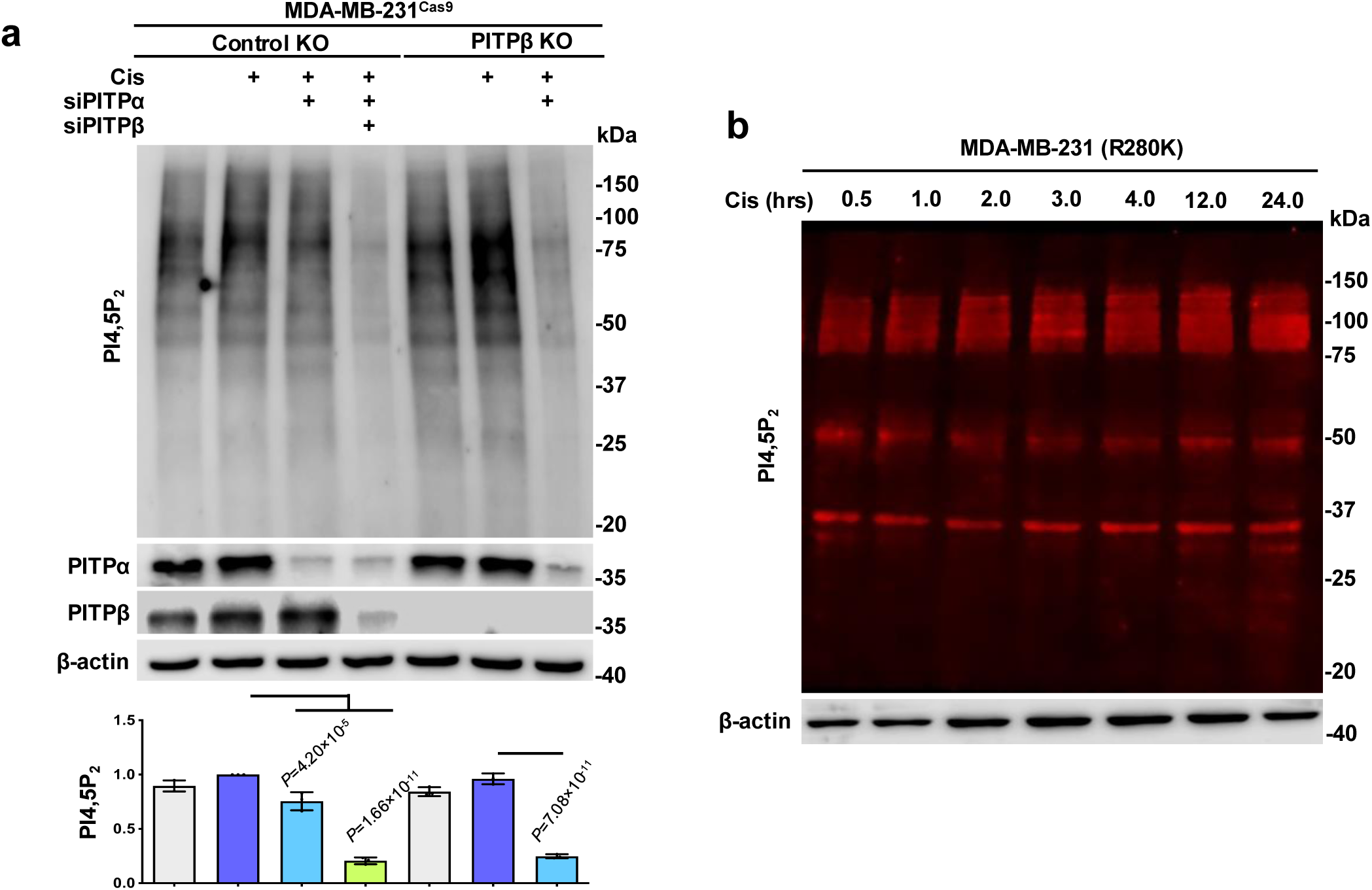
PITPα/β regulate stress-responsive, unique, and stable protein-PIP_n_ complexes. **a**, MDA-MB-231^Cas9^ cells with PITPβ KO and control non-targeted KO were transfected with control siRNAs or siRNAs against PITPα or both PITPα and PITPβ. After 24 h, cells were treated with 30 µM cisplatin or vehicle for 24 h before being processed for WB against PI4,5P_2_ and quantified by ImageJ. n=3 independent experiments. **b**, MDA-MB-231 cells were treated with 30 μM cisplatin and processed for WB against PI4,5P_2_ at the indicated treatment timepoints. n=3 independent experiments. For all graphs, data are presented as the mean ± SD.

**Extended Data Figure 7.**
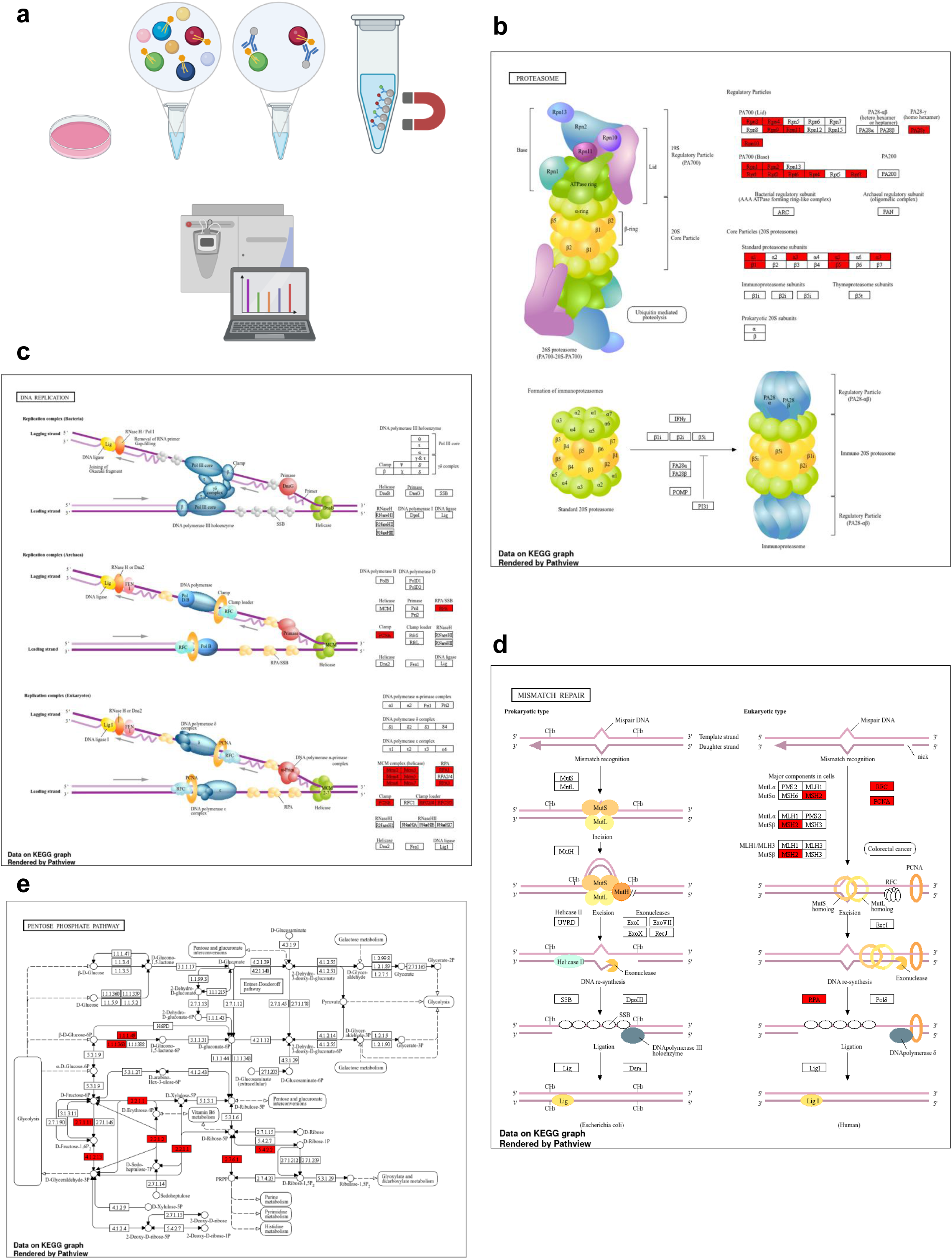
Protocol and pathway diagrams from proteomic analysis. **a**, Graphical diagram depicting IP-MS approach using an anti-PI4,5P_2_ antibody and Dynabeads to enrich for protein-PI4,5P_2_ complexes. **b-e**, Pathview rendering of enriched pathways identified using IP-MS of protein-PI4,5P_2_ complexes with specific targets highlighted in red. Redering produced by ShinyGO 0.85 and KEGG pathway definitions for proteosome (**b**), DNA replication (**c**), Mismatch repair (**d**), and pentose phosphate pathway (**e**).

